# rapidPop: Rapid population assessments of wildlife using camera trap data in R Shiny Applications

**DOI:** 10.1101/2020.03.30.017103

**Authors:** Michael A. Tabak, Jesse S. Lewis, Peter E Schlichting, Nathan P. Snow, Kurt C. VerCauteren, Ryan S. Miller

## Abstract

rapidPop is a new R package for implementing occupancy models for Rapid Population Assessments (RPAs) with data from camera traps. RPAs are designed to provide quick assessments of a density index so that users can identify relative changes in density associated with changes in conditions (Schlichting *et al.*, 2020). For example, users may want to assess if there was a change in density after an effort to cull an invasive species. rapidPop provides a Shiny Application for running occupancy models with the option to include the effect of parameters on occupancy (e.g., before or after a culling operation).

## Description

rapidPop is an R package for implementing occupancy models from data collected in wildlife Rapid Population Assessments (RPAs). This method is described in detail by Schlichting *et al.* (2020). This package implements occupancy models using the R package unmarked (Fiske & Chandler, 2011). The R packages shiny v. 1.4.0.2 (Chang *et al.*, 2019) and shinyFiles v. 0.7.5 (Pedersen *et al.*, 2019) are used to create Shiny Applications (‘Shiny Apps’) in rapidPop, which facilitate a graphical user interface (GUI), an environment where users can manually select their input files and run occupancy models. rapidPop was built using R 3.6.3 (R Core Team, 2020).

The model structure implemented in rapidPop is a zero-inflated binomial where the latent, un-observed occupancy status of site *i* (*Z_i_*) depends on the probability of occupancy at that site (*ψ_i_*):

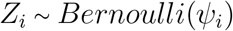

The observation process is modeled by:

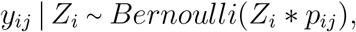

where *y_ij_* are the observations (0 indicating no individuals observed and 1 indicating ≥ 1 observed individual) at site *i* in occurrence *j* and *p_ij_* is the ‘detection probability,’ or the probability of detecting an individual given that one is present (Mackenzie, 2006; Royle & Dorazio, 2008).

rapidPop offers the option to model the effect of a parameter on occupancy. If this option is employed, the model structure includes a process model, where occupancy probability is modeled as a function of the parameter (X):

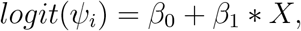

where β_1_ represents the effect of the parameter on occupancy.

## Usage

The R package rapidPop is freely available from GitHub. Users can download and install the software by typing `devtools::install_github(“mikeyEcology/rapidPop”)` into the R console. To launch the Shiny App that will conduct occupancy modeling, type `rapidPop::runShiny(“occMod”)` and a Shiny App will open. To select your file containing observations, click on the button `Input file` (see Fig. 1); you can then navigate to your file. This file can be formatted either as `.txt or.csv.` You will need to specify the file type in the next option. The file must contain columns with observations of the animals. Enter the column headers (column names) of the first column containing observation data and the final column containing observation data in the appropriate boxes. For example, if your observation data were contained in columns named `day1, day2,…, day12`, The first column would be `day1` and the final column would be `day12`. This type of naming strategy is reflected in the example input file that is included in rapidPop and implemented in Figure 1.

**Figure 1:**
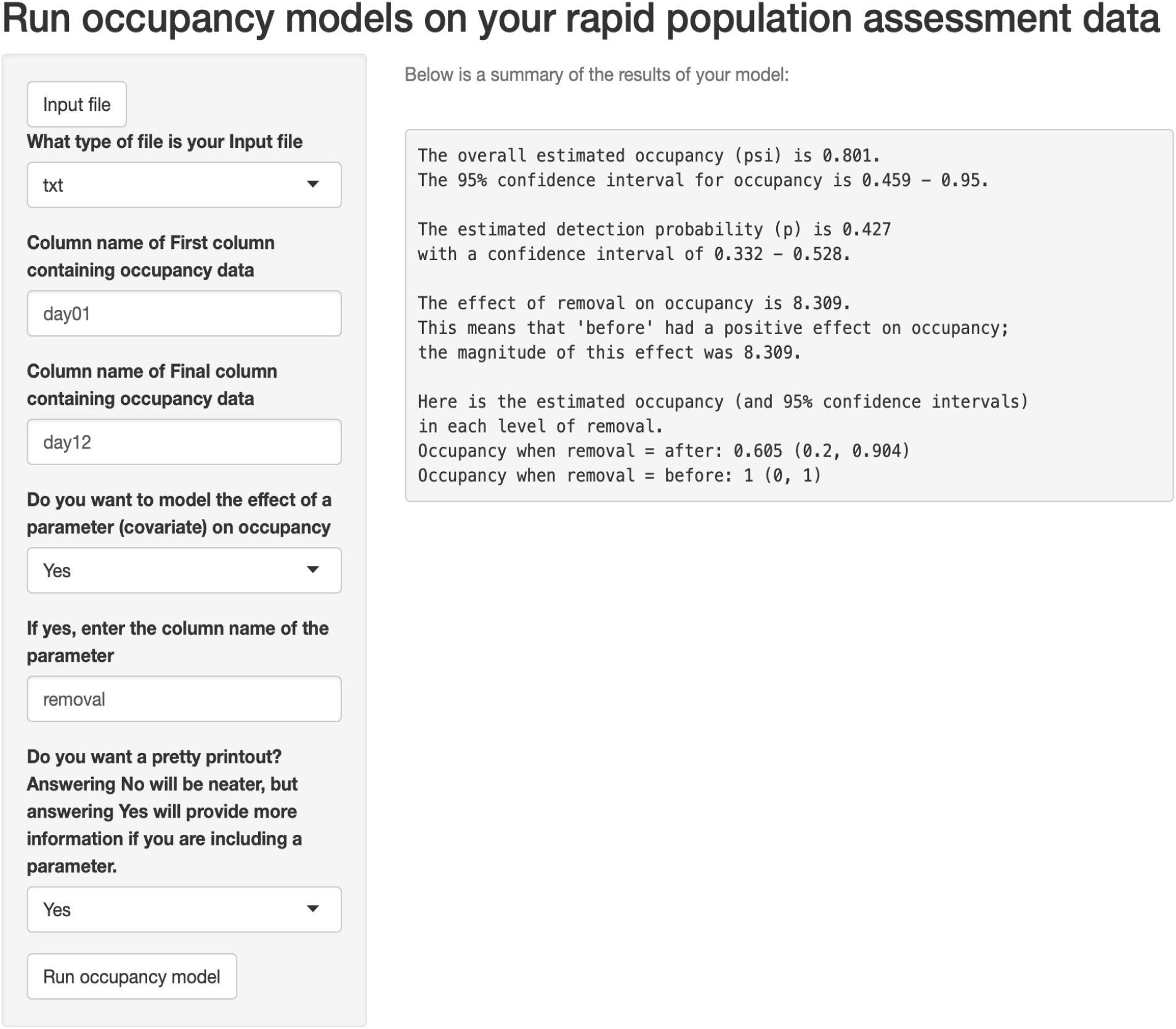
Example of the Shiny App, including the informational summary of the model results

rapidPop provides the option to model the effect of a parameter on occupancy. If you would like to do this, select `Yes` from this box and specify the column name of the parameter in your input file. For example, a user might want to model the effect of a management intervention on occupancy. If this was removal, you could have a column in your input file called `removal` with values `before` and `after`. You would then enter `removal` as the parameter and the model will show the effect of removal. By default, this application will print out the results in a summarized format; if you prefer more details, select No for the final option. For more details about the theory and methods of RPAs, see Schlichting *et al.* (2020).

Note that the `devtools` package, which is required for installing the R packages from GitHub usually requires an installation of the Rtools software on Windows computers. Rtools is freely available for download from CRAN (https://cran.r-project.org/bin/windows/Rtools/).

## Example

After installing rapidPop and typing `rapidPop::runShiny(“occMod”)` into the R console, a GUI window will open (e.g., Fig. 1). In the example illustrated by Figure 1, the estimated occupancy (ψ; the probability that a given site is occupied) is 0.801, with a 95% confidence interval around *ψ* of 0.459 - 0.950, indicating that about 4/5 sites are occupied, but there is quite a bit of uncertainty around this estimate. The detection probability (p; the probability of detecting an individual in sampling given that one is present) is 0.427 with a 95% confidence interval of 0.332 - 0.528. This means that an individual is detected in sampling (e.g., found in a camera trap image) about half of the time that one is present. The *β_1_* value for effect of removal on occupancy is 8.309. The next line shows the direction of this effect, specifically, ‘before’ has a positive effect on occupancy, meaning that occupancy decreases after removal. Finally, the occupancy estimates under each condition are shown. Occupancy before removal was 1 (each site was occupied), and after removal occupancy was 0.6 (3/5 sites were occupied). The example data file that produced the results in Figure 1 is included with rapidPop.

